# Regulatory involvement of the PerR and SloR metalloregulators in the *Streptococcus mutans* oxidative stress response

**DOI:** 10.1101/2020.12.15.422995

**Authors:** Talia R. Ruxin, Julia A. Schwartzman, Cleo R. Davidowitz, Robet A. Haney, Grace A. Spatafora

**Affiliations:** Department of Biology, Middlebury College, Middlebury, Vermont, USA; Department of Civil and Environmental Engineering, Massachusetts Institute of Technology, Massachusetts, USA; Department of Biology, Ball State University, Muncie, Indiana, USA

## Abstract

*Streptococcus mutans* is a commensal of the human oral microbiome that can instigate dental caries under conditions of dysbiosis. This study investigates *S. mutans* metalloregulators and their involvement in mediating a response to oxidative stress. Oxidative stress in the oral cavity can derive from temporal increases in reactive oxygen species (ROS) after meal consumption, and from endogenous bacterial ROS-producers that colonize the dentition as constituents of dental plaque. We hypothesize that the PerR (SMU.593) and SloR (SMU.186) metalloregulatory proteins in *S. mutans* contribute to oxidative stress tolerance by regulating the expression of genes responsive to H_2_O_2_ challenge. The results of qRT-PCR experiments with *S. mutans* cultures exposed to 0.5mM H_2_O_2_ reveal *perR* transcription that is responsive to the peroxide stressor, and *sloR* transcription that is subject to PerR repression. The results of gel shift assays support direct binding of a PerR homolog to the *S. mutans sloR* promoter at Fur and PerR consensus sequences on the UA159 chromosome. In addition, transcription of the *S. mutans tpx* and *dpr* antioxidant genes is upregulated in a *perR/sloR* double knockout mutant, consistent with heightened resistance of the *S. mutans* GMS802 *perR*-deficient strain when challenged with H_2_O_2_. Cumulatively, these results reveal a relationship of reciprocity between the PerR and SloR metalloregulators during the *S. mutans* response to oxidative stress and begin to elucidate the fitness strategies that evolved to foster *S. mutans* survival and persistence in the transient environments of the human oral cavity.

**IMPORTANCE:** In 2020, untreated dental caries, especially in the permanent dentition, ranked among the most prevalent infectious diseases worldwide. Moreover, caries disproportionately affects children and individuals of low socioeconomic status. Untreated caries can lead to systemic health problems and has been associated with extended school and work absences, inappropriate use of emergency departments, and an inability for military forces to deploy. In combination with public health policy, research aimed at alleviating *S. mutans-induced* tooth decay is important because it can improve oral health, as well as overall health, especially for underserved populations. This research is focused on the *S. mutans* SloR and PerR metalloregulatory proteins that can help inform the development of therapeutics aimed at alleviating and potentially preventing dental caries.

## INTRODUCTION

Dental caries is the most common chronic infectious disease in the US and around the world, affecting approximately 60 to 90% of school-age children and a vast majority of adults (1). As of 2020, untreated dental caries in the permanent dentition ranked as the most highly prevalent infectious disease worldwide (2). Maladies of the oral cavity, including dental caries, are also inextricably linked to systemic diseases such as diabetes, cardiovascular disease, and low birth weights (3). Moreover, the social determinants of health widens the gap between those with access to quality oral healthcare and under resourced populations, namely minority racial and ethnic groups and those of low socioeconomic status (4, 5).

*Streptococcus mutans*, one of the primary causative agents of dental caries in humans (6), uses dietary sucrose as a substrate for extracellular polysaccharide synthesis, facilitating formation of the plaque biofilm. Subsequent acid production demineralizes tooth enamel which can lead to decay (7). In addition to its unique aciduric and acidogenic properties that allow *S. mutans* to withstand stressors such as acid pH are virulence factors that protect it against oxidative stress, both during and between mealtimes (8–10). Especially during mealtimes, reactive oxygen species (ROS) can reach high levels in the plaque environment that extraoral microbes are not equipped to detoxify. In contrast, *S. mutans* has evolved an oxidative stress response that can detoxify ROS that derive from exogenous sources, as well as from within the dental plaque microbiome where the H_2_O_2_-producing *Streptococcus sanguinis* and *Streptococcus gordonii* reside (11, 12). H_2_O_2_ production evolved in these early colonizers of the plaque biofilm as a fitness strategy aimed at antagonizing neighboring microbes, such as the non-peroxigenic *S. mutans* which are later colonizers of the dentition. As a constituent of the plaque microbiome however, *S. mutans*, can prompt the fine-tuning of its metal ion uptake machinery, thereby curtailing ROS production and toxicity (13–15). Taken together, the overall fitness of *S. mutans* in the oral cavity is highly dependent on the expression of virulence factors that allow it to coordinate an appropriate response to ROS- (and more specifically, peroxide-) induced stress.

Bacterial metalloregulatory proteins are the “watchdogs” of intracellular metal ion homeostasis; they are essential for *S. mutans* survival and pathogenesis. Among the metalloregulatory proteins in *S. mutans* is PerR, an 18kDa Fur family regulator that functions as a metal ion-dependent transcriptional repressor (16, 17). PerR is the sole Fur family regulator in *S. mutans*, consistent with an *in silico* analysis of the *S. mutans* UA159 chromosome (homd.org) which reveals a single 477-bp *perR* gene (SMU.593). Multiple Fur homologs, including Fur and PerR, have been documented in other bacteria, such as *Bacillus subtilis* (16). While little is known about PerR in *S. mutans*, a recent report describes the transcriptomes of a *perR* insertion-deletion mutant and its wild type progenitor in which a relatively small number of genes are differentially expressed (18, 19). In contrast, a 25kDa SloR protein belonging to the DtxR family of metalloregulators modulates a plethora of *S. mutans* genes, including those that encode essential metal ion transport (20–22). Specifically, *S. mutans* SloR regulates an ABC-type metal ion transport operon comprised of the *sloABC* genes, by mechanisms that have been previously described (23, 9, 24, 22). A more recent report describes an ancillary metal ion transporter, called MntH, that helps maintain intracellular metal ion homeostasis in *S. mutans*, and which bears similarity to Nramp-type transporters (25). Taken together, the SloABC and MntH transporters as well as their regulators are essential for ensuring *S. mutans* fitness and survival in both healthy and dysbiotic plaque environments (9, 25, 26).

In the present study, we set out to understand the mechanism(s) that allow *S. mutans* to tolerate oxidative stress in the plaque microbiome. Herein, we demonstrate that *perR*-deficient strains of *S. mutans* are more resistant to hydrogen peroxide stress than their wild type progenitors, consistent with previous reports (18, 27), and that PerR is a direct repressor of *sloR* gene transcription when challenged with peroxide stress. In addition, we describe a relationship between the PerR and SloR metalloregulatory proteins during the *S. mutans* response to peroxide stress, in part by confirming their roles in controlling expression of the *S. mutans* oxidative stress response genes, *tpx* and *dpr*.

## RESULTS

The *S. mutans* GMS802 *perR* insertion-deletion mutant is more resistant to peroxide challenge than its UA159 wild type progenitor. To determine if disruption of the *perR* gene impacts the *S. mutans* peroxide stress response, we grew the *S. mutans* UA159 wild type strain and its isogenic *perR* insertion-deletion mutant, GMS802, in THYE broth supplemented with a sublethal concentration of H_2_O_2_ (0.0015%). The results of these growth determination assays revealed a generation time for GMS802 that was significantly shorter than that of the UA159 wild type strain (50.5 ± 0.7 min versus 54.4 ± 1.3 min, respectively, p<0.05) in the presence of the peroxide stressor. To mimic conditions of peroxide stress *in vivo*, we repeated these experiments in a conditioned medium containing *S. gordonii*-generated H_2_O_2_ where we likewise noted a significant decrease (p<0.05) in GMS802 generation time (52.3 ±1.1 min) relative to that of UA159 (56.5 ± 1.6 min). Collectively, these data indicate that growth of the GMS802 *perR* mutant is more rapid than wild type UA159 when challenged with the peroxide stressor at physiological concentrations. In contrast, we noted no significant difference in generation time between UA159 (44.5± 1.0 min) and GMS802 (43.2 ± 0.8 min) when both strains were grown in the absence of H_2_O_2_-challenge.

To determine if the *perR* insertion-deletion mutation promotes *S. mutan*s survival when grown adjacent to peroxide-producing *S. gordonii*, we performed *S. gordonii* antagonism assays as previously described (12). The results of these assays reveal inhibition of the *S. mutans* UA159 wild type strain when cultivated adjacent to *S. gordonii* cells that had been growing and producing H_2_O_2_ for 16 hours (Fig. 1A). In contrast, the GMS802 *perR* mutant appeared to be unaffected by *S. gordonii* peroxide challenge (Fig. 1B). In fact, the mutant appeared to infiltrate the growth zone of *S. gordonii* despite the elaboration of H_2_O_2_, consistent with a phenotype of peroxide resistance. Complementation of the *perR* mutation *in cis* in the GMS803 strain restored peroxide sensitivity to near wild type levels (Fig. 1C).

**Fig. 1.**
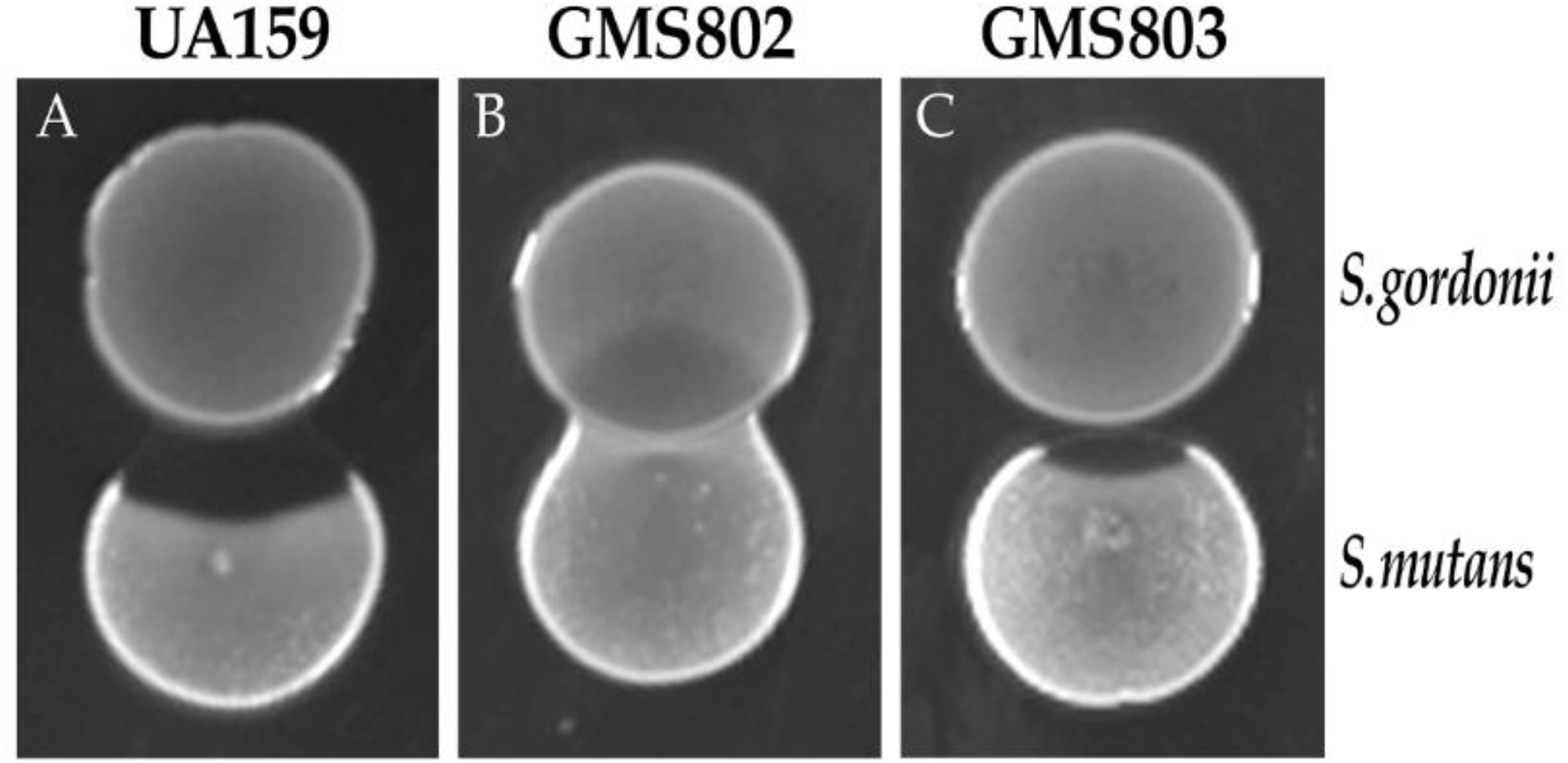
The *S. mutans* GMS802 mutant is more resistant to hydrogen peroxide production by *S. gordonii* than its UA159 wild type progenitor. Peroxigenic *S. gordonii* was spot inoculated onto the surface of a THYE agar medium and grown for 16 hours before inoculating the *S. mutans* UA159 wild type strain in close juxtaposition. **A.** Following overnight incubation at 37°C and 5% CO_2_, inhibition of *S. mutans* growth is evident. **B.** Growth of the *perR*-deficient GMS802 mutant was not inhibited by *S. gordonii* peroxide challenge under these same growth conditions. **C.** Complementation of the *perR* mutation in *S. mutans* GMS803 restored its sensitivity to peroxide challenge by *S. gordonii*. Each panel represents a single representative experiment among the eight total biological replicates performed.

### Peroxide stress cues transcription of the *S. mutans perR* gene, but not *sloR*

Reports in the literature describe PerR as a redox sensor in *Bacillus subtilis* and *Streptococcus pyogenes* (28, 29). Accordingly, we hypothesized that oxidative stress would induce *perR* gene transcription in *S. mutans*. To test this hypothesis, we challenged *S. mutans* wild type UA159 cultures with 0.5mM H_2_O_2_ and monitored *perR* expression in these and unchallenged controls in qRT-PCR experiments. The experimental results indicate that *perR* transcription was upregulated 2.5-fold in *S. mutans* UA159 cultures challenged with H_2_O_2_ compared to unchallenged cultures (n=4, p<0.05; Fig. 2A). In contrast, exposure of *S. mutans* UA159 to the H_2_O_2_ stressor did not have any impact on *sloR* gene transcription relative to controls (n=5; p>0.05; Fig. 2B).

**Fig. 2.**
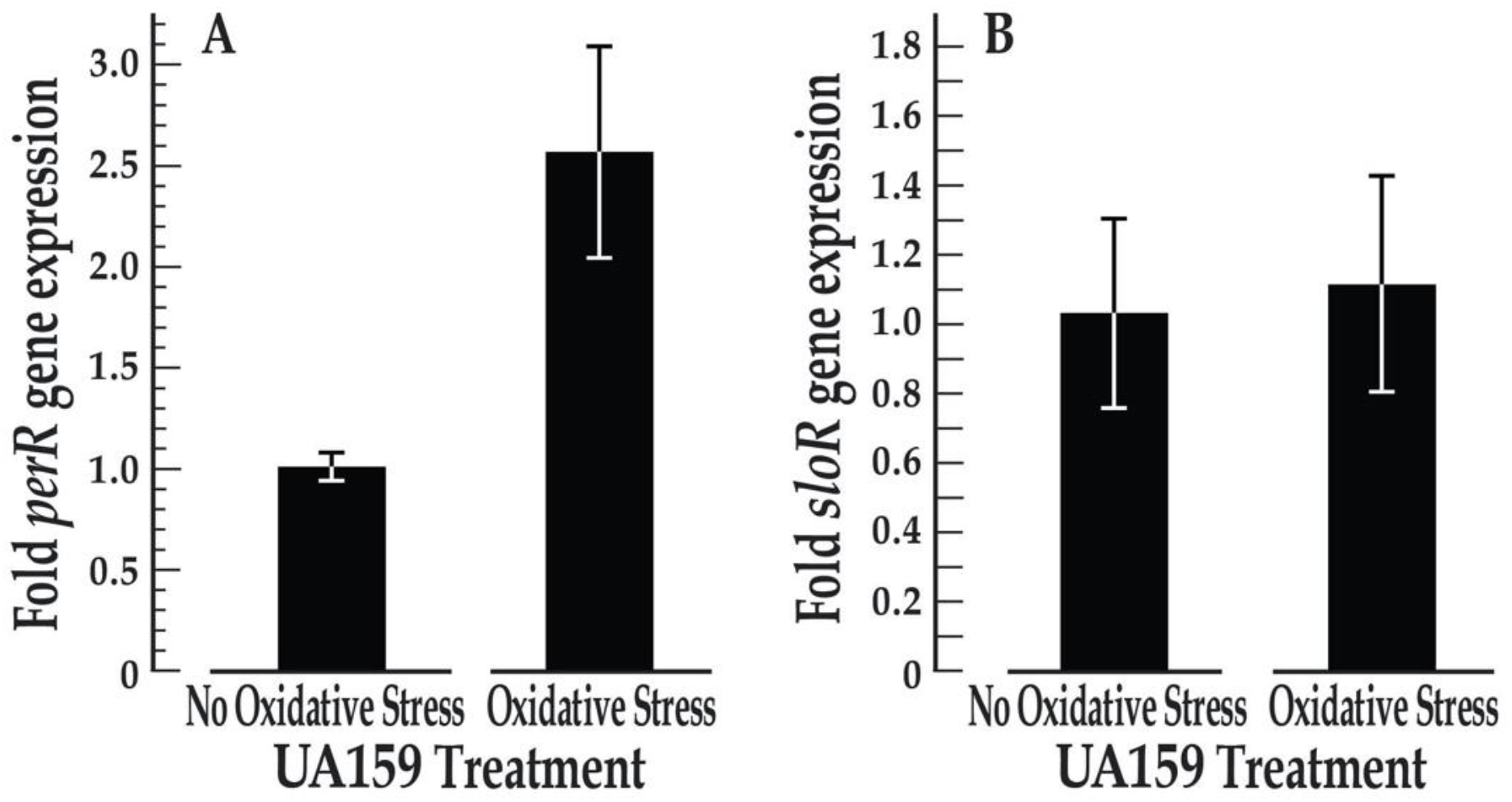
Peroxide challenge increases transcription of the *perR* gene, but not the *sloR* gene, in *S. mutans* UA159. Expression of the *S. mutans perR* and *sloR* genes was determined in qRT-PCR experiments and normalized against an endogenous *hk11* control gene. **A.** Expression of *S. mutans perR* is increased 2.5-fold in wild type UA159 cultures grown in the presence of 0.5mM H_2_O_2_ (n=4, p<0.05). **B.** Transcription of the *S. mutans* UA159 *sloR* gene is not significantly impacted under the same test conditions (n=4, p>0.05). Error bars depict SEM.

### *In silico* prediction of PerR and Fur binding sites in *S. mutans*

Since *S. mutans* contains only one Fur family metalloregulator with an unknown binding motif, we used a combination of consensus binding weight matrices based on known PerR and Fur binding motifs in other bacteria to identify targets of *S. mutans* PerR. Specifically, we used four consensus binding weight matrices based on the binding sites of PerR homologs in *S. pyogenes* and *S. suis* (consensus: TTAGAATCATTWTAA), PerR (consensus AWTTAKAATAATTATAATT) and Fur (consensus: NWAAATGATAATCATTATCAWT) homolog binding sites in *Bacillus subtilis, Staphylococcus aureus*, and the *Streptococcus* species, and Zur, Fur and PerR, homolog binding sites (consensus TNWWAWTKATAATMATTATMAWTTAN) in *B. subtilis, Listeria monocytogenes, B. amyloliquefaciens, S. aureus, S. pyogenes* and *S. suis*. All four matrices were used to screen 1113 *S. mutans* intergenic regions with a length of at least 50-bp. We identified a total of 3824 significant (p-value <0.001, q-value < 0.129) motif matches with a boundary localized to the putative promoter within 50-bp of the start codon. After purging redundancies by retaining only the match with the lowest p-value for a given promoter region in a given matrix search, we identified a total of 779 genes with a match from one or more matrices in the adjacent promoter region (see supplementary Dataset S1). A total of 191 gene promoter regions revealed binding motif matches with all four matrices, including those that precede the *S. mutans dpr* (SMU.540), *ahpC*, (SMU.764) and *gor* (SMU.838) antioxidant genes, as well as the *S. mutans sloR* gene (SMU.186; Fig. 3).

**Fig. 3.**
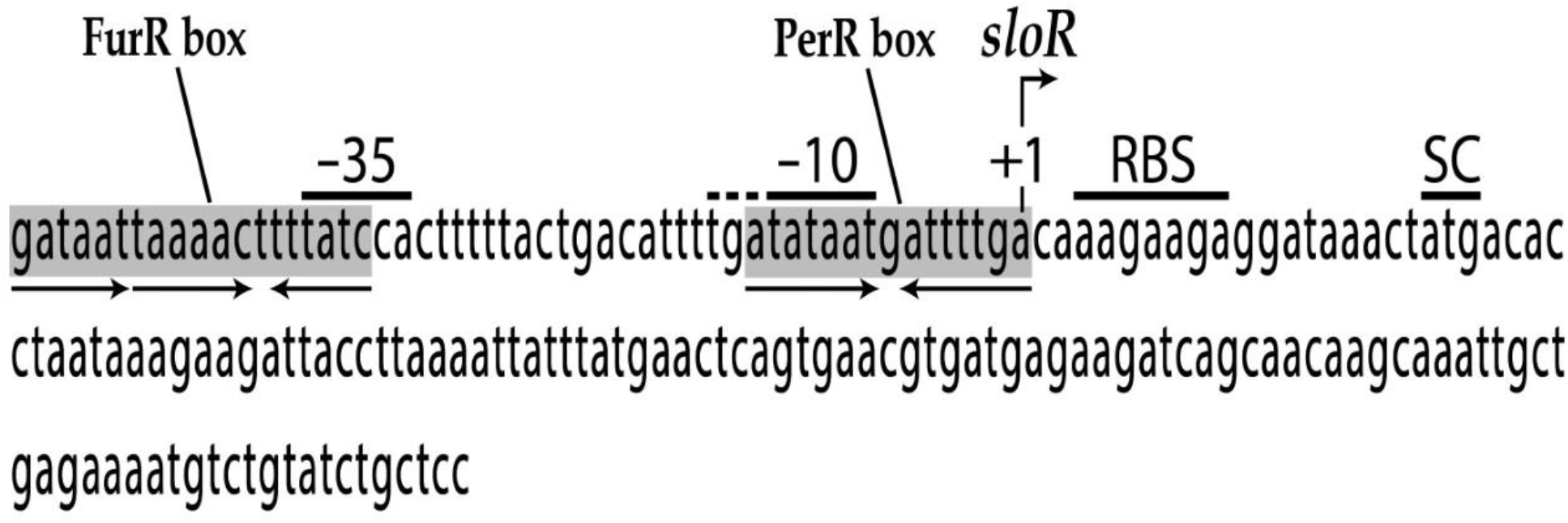
The *sloR* promoter includes putative consensus binding sequences for the *S. mutans* PerR metalloregulator. Shown is the intergenic region between the *S. mutans sloC* and *sloR* genes on the *S. mutans* UA159 chromosome. The nucleotide sequence in this region was aligned with the *S. mutans* UA159 genome from the NCBI GenBank database (RefSeq accession number NC_004350.2). The sequence harbors the –35 and extended –10 promoter regions that drive *sloR* gene transcription from the +1 transcription start site (designated by the bent arrow above the sequence). Also shown is the predicted ribosome binding site (RBS) and the start codon (SC) for the *sloR* coding sequence. *In silico* analysis of the *sloR* promoter supports the presence of a Fur binding sequence in the –35 region, comprised of two forward hexamers separated by 1-bp from a third inverted hexamer (6-6-1-6), and a predicted 7-1-7 PerR binding sequence that shares overlap with the –10 promoter region (gray shading).

### PerR is a repressor of *sloR* gene transcription

To reveal whether *S. mutans* PerR modulates expression of *sloR* under conditions of H_2_O_2_ stress, qRT-PCR experiments were performed with GMS802 and UA159 cultures that had been exposed to peroxide challenge. We observed a 2-fold increase in *sloR* transcription in GMS802 compared to UA159 when these *S. mutans* strains were challenged with 0.5mM H_2_O_2_, indicating that PerR functions as a repressor of *sloR* transcription in *S. mutans* wild-type cells grown under conditions of oxidative stress (n=4, p<0.05; Fig. 4A). When the *S. mutans* GMS584 *sloR*-deficient strain was grown under the same test conditions, expression of *perR* was not significantly impacted, indicating that *perR* transcription is not subject to control by the SloR metalloregulator (n=5, p>0.05, Fig. 4B).

**Fig. 4.**
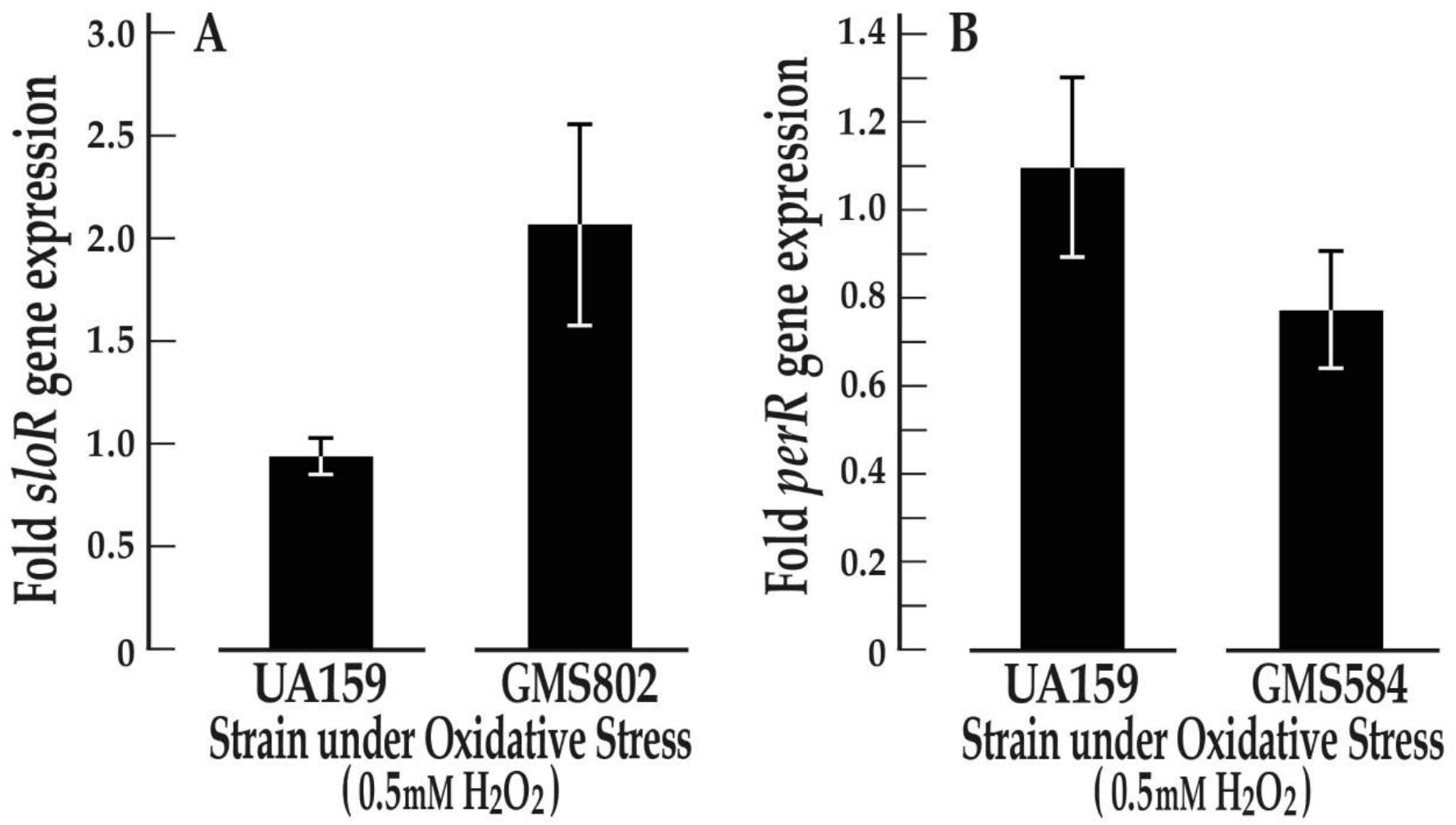
Transcription of the *S. mutans sloR* and *perR* genes in the presence of 0.5mM H_2_O_2_. **A.** Expression of the *S. mutans sloR* gene is 2-fold greater in the GMS802 *perR* insertion-deletion mutant compared to the wild type UA159 strain (n=4, p<0.05) when challenged with 0.5mM H_2_O_2_ for 15 minutes. **B.** Expression of the *S. mutans perR* gene is not significantly different in the GMS584 *sloR* insertion-deletion mutant compared to wild-type (n=5, p>0.05). Expression was normalized against that of an endogenous *hk11* control gene. Error bars depict SEM.

### A PerR homolog from *Bacillus subtilis* binds directly to the *S. mutans sloR* promoter

To elucidate whether the impact of PerR on *sloR* transcription is direct, we performed gel mobility shift assays with a 17.4kDa PerR homolog in *B. subtilis*, called Fur, and a *sloR* promoter target probe that harbors putative Fur- and Per-like consensus binding sequences as predicted by the *in silico* analysis described above. The outcome of experiments performed with the PerR homolog support direct Fur binding to the 53-bp *sloR* promoter probe at protein concentrations as low as 60nM (Fig. 5A). The addition of EDTA to a reaction mixture containing as much as 400nM Fur protein abrogated the band shift, consistent with a Fur-DNA interaction that is metal ion-dependent.

**Fig. 5.**
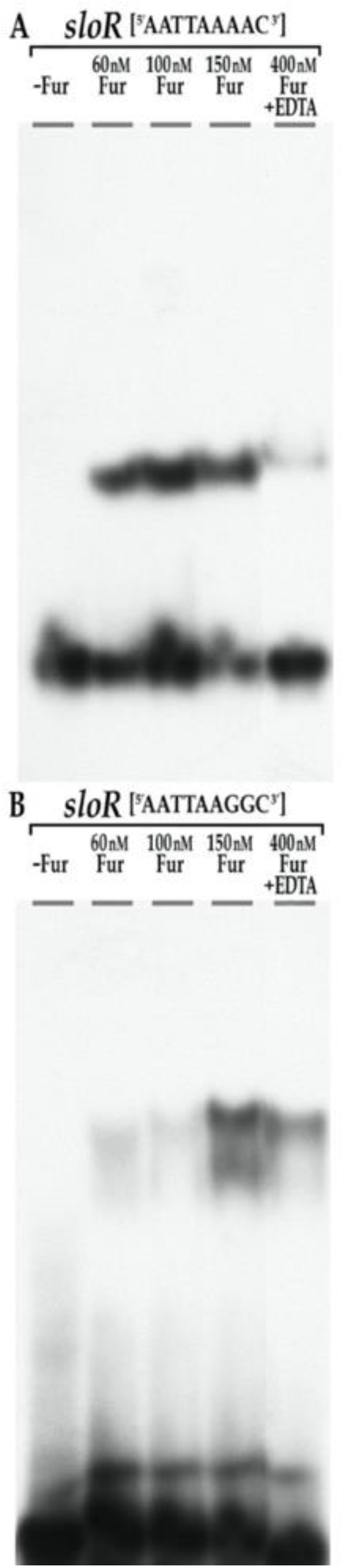
A PerR homolog from *B. subtilis* shifts a sequence within the *S. mutans sloR* promoter region in gel mobility shift experiments. A 17.4kDa Fur protein from *Bacillus subtilis* (kind gift of Dr. John Helmann, Cornell University) was used in gel mobility shift experiments with a 53-bp DNA amplicon that harbors the wild-type *S. mutans sloR* promoter including regions –35 and –10 **(A)** or a 54-bp *sloR* promoter sequence that is degenerate within the predicted Fur binding sequence that shares overlap with the –35 region **(B)**. Reaction mixtures were resolved on a 12% non-denaturing polyacrylamide gel for 450 volt hours. **A.** As little as 60nM Fur generated a robust shift of the *sloR* target sequence. **B.** Shifting of the 54-bp degenerate *sloR* target sequence was compromised even when as much as 100nM Fur was added to the reaction mixture. Two bands are evident when 150nM Fur was included in the reaction mixture, consistent with two separate binding sites on the *sloR* target probe. Addition of EDTA abrogated the band shifts, consistent with a Fur-DNA interaction that is metal-ion-dependent.

### Binding of a PerR homolog from *Bacillus subtilis* to a degenerate *sloR* promoter in *S. mutans* is compromised

To validate sequence-specific binding of the heterologous Fur protein from *B. subtilis* to the *S. mutans sloR* promoter, we performed gel shift assays with a degenerate 54-bp oligonucleotide containing two A to G transition mutations in the predicted Fur-consensus sequence that overlaps the −35 region of the *sloR* promoter (AATTAA**GG**C). The experimental results indicate that *in vitro* binding of 60nM Fur to the degenerate *sloR* target sequence was compromised when compared to the robust binding we observed when 60nM Fur was allowed to react with the wild-type target sequence (mean pixel area for the band shift with the degenerate probe was 63% versus 83% with the wild type probe) (Fig. 5B). Moreover, robust binding to the degenerate sequence required at least 150nM Fur protein, consistent with an attenuated Fur-DNA interaction, and an important role for adenine nucleotides in accommodating the binding of Fur to the wild-type consensus sequence in *S. mutans*.

### *S. mutans* PerR and SloR repress *dpr* and *tpx* transcription

We hypothesized that the heightened peroxide-resistance we observed for the GMS802 *perR*-deficient strain might reflect repression of the wild type antioxidant arsenal in UA159. To explore the putative impact of the PerR and SloR metalloregulators on two of these antioxidant genes, *dpr*, and *tpx*, we performed qRT-PCR experiments with RNAs isolated from H_2_O_2_-challenged UA159 (wild type), GMS802 (*perR*-deficient), GMS584 *(sloR*-deficient), and GMS1386 (*perR*- and *sloR*-deficient) *S. mutans* strains. Notably, transcription of *dpr* and *tpx* was highly variable in the single mutants compared to wild type (data not shown) but was clearly repressed in the double knockout mutant. Specifically, expression of the *dpr* and *tpx* genes in *S. mutans* GMS1386 was significantly increased up to 3-fold when compared with that in the UA159 wild type strain (Fig. 6A and 6B). Thus, under conditions of peroxide stress it seems that PerR and SloR work together to downregulate transcription of *S. mutans tpx, dpr*, and likely other antioxidant genes.

**Fig. 6.**
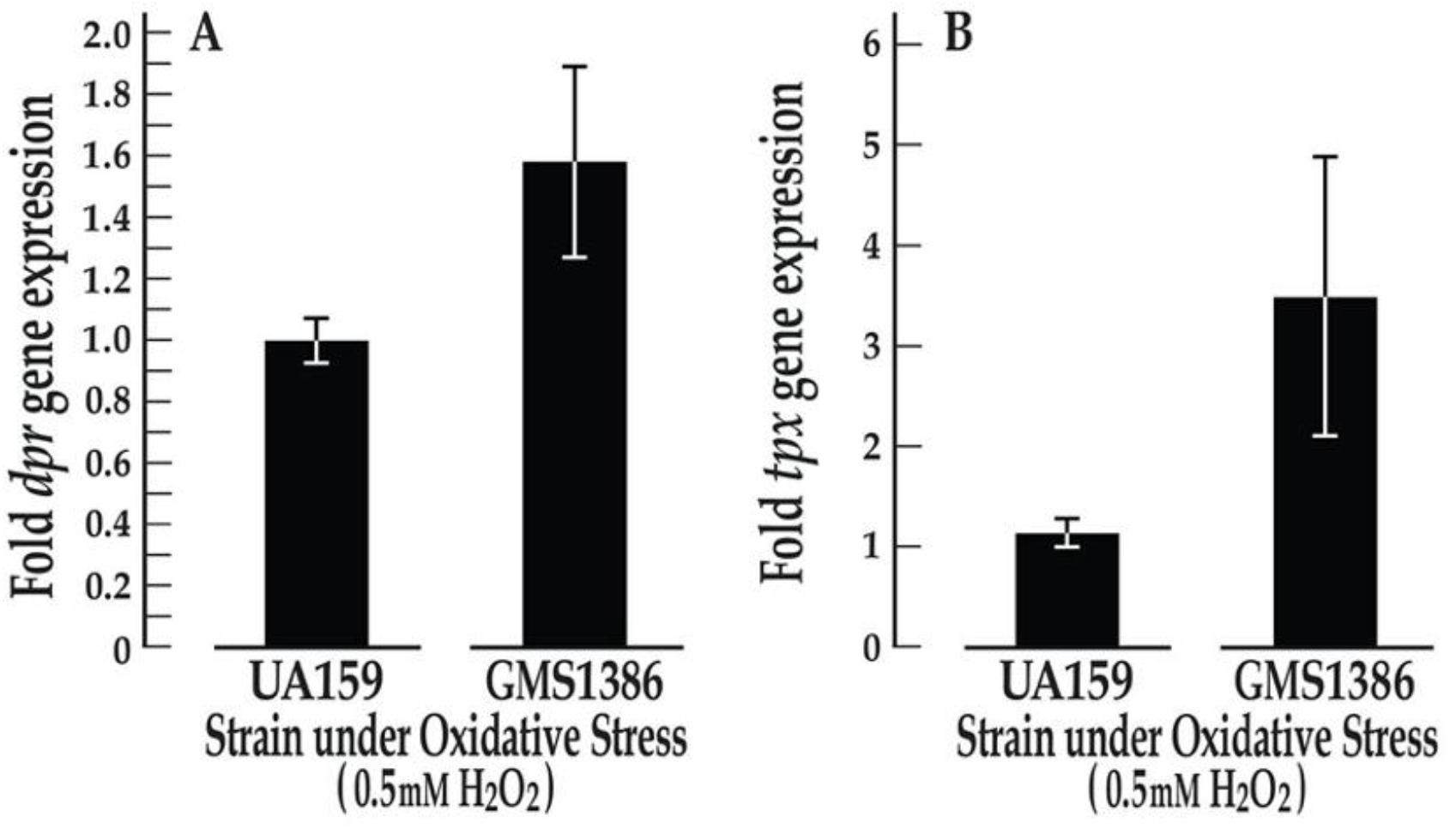
*S. mutans dpr* and *tpx* gene transcription is upregulated in the *S. mutans* GMS1386 PerR/SloR double mutant compared to the UA159 wild type strain when challenged with 0.5mM H_2_O_2_. **A.** *S. mutans dpr* expression is increased 1.6-fold in the *S. mutans* GMS1386 knockout mutant compared to its UA159 wild type progenitor under conditions of peroxide stress (n=5, p<0.05). **B.** Transcription of the *S. mutans tpx* gene is heightened >3-fold in GMS1386 compared to UA159 in the presence of peroxide stress (n=3, p<0.05). Expression of the *dpr* and *tpx* genes was normalized to that of an endogenous *hk11* control gene. Error bars depict SEM.

### Regulation of *tpx* transcription by PerR is not direct

We performed gel mobility shift experiments with *S. mutans* UA159 and GMS802 whole cell lysates (WCL) and a 242bp target probe that harbors the *tpx* promoter to reveal whether the impact of PerR on *tpx* transcription is direct. The experimental results reveal no evidence of shifting the *tpx* promoter region even when as much as 20ug of protein lysate was included in the reaction mixture (Fig. 7). The *sloR* promoter was used as a positive control in these experiments.

**Fig. 7.**
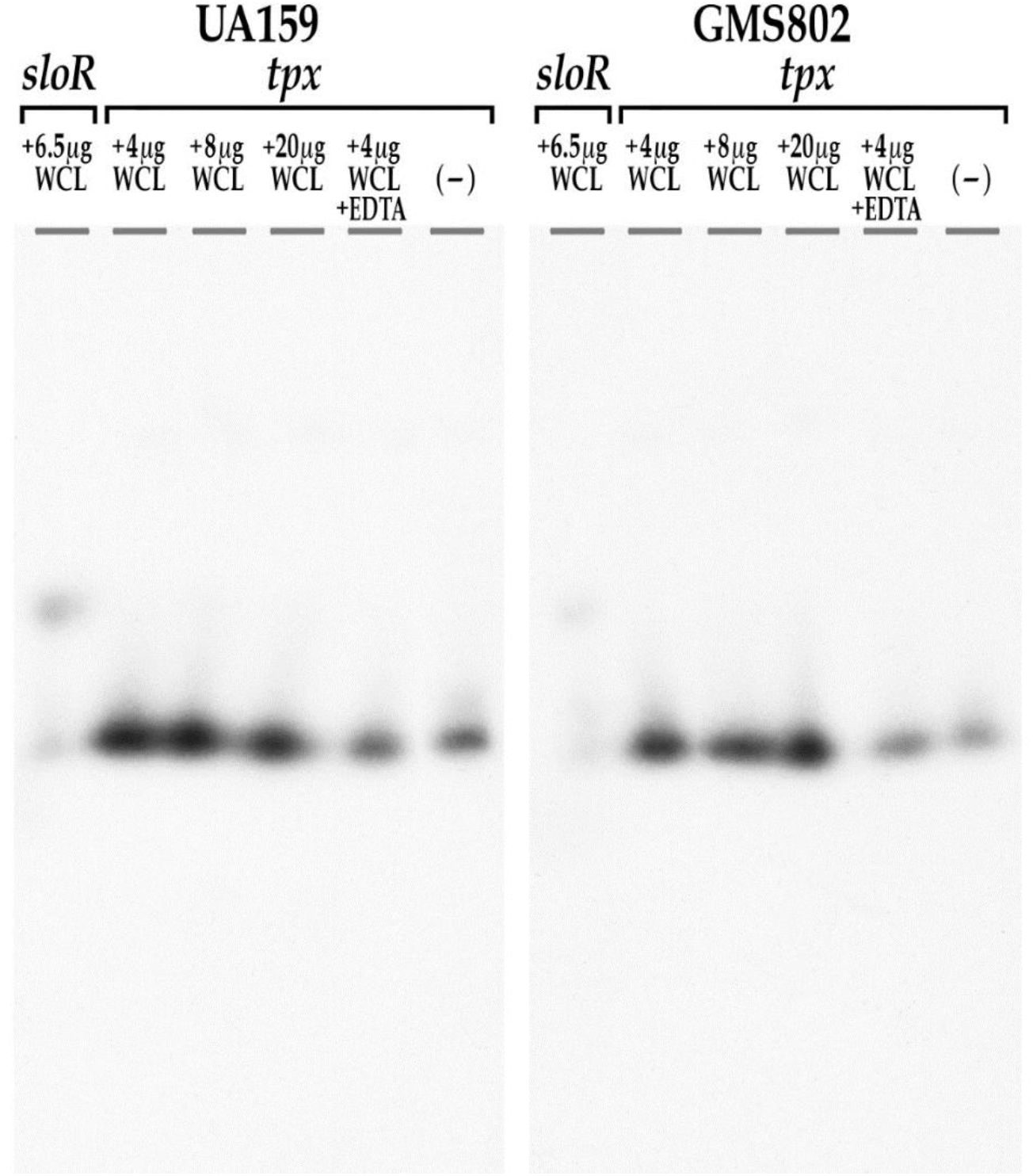
Whole cell lysates fail to shift the *S. mutans tpx* promoter in gel shift experiments. Whole cell lysate prepared from *S. mutans* UA159 and GMS802 failed to shift a 242-bp amplicon that harbors the *S. mutans tpx* gene promoter. The 53-bp *sloR* promoter was used as a positive control. [γ-^32^P] end-labeled DNA amplicons were resolved on a 12% non-denaturing polyacrylamide gel for 225 volt hours.

## DISCUSSION

In the present study, we describe a GMS802 *perR*-deficient strain of *S. mutans* that is more resistant to hydrogen peroxide challenge than its wild type UA159 progenitor, corroborating other published reports of peroxide resistance in PerR-deficient strains of *S. mutans* (18, 27). Yet other reports in the literature (20) describe a SloR-deficient GMS584 strain of *S. mutans* that demonstrates compromised survivorship in the face of peroxide challenge. Taken together, these findings support a role for both PerR and SloR in *S. mutans* aerotolerance. Herein, we extend these reported findings by characterizing the relationship between the PerR and SloR metalloregulators in the *S. mutans* response to H_2_O_2_ challenge, and by elucidating PerR binding to the *S. mutans sloR* promoter region that harbor Fur- and PerR-like consensus binding sequences.

Not surprisingly, we describe *perR* transcription that is upregulated in *S. mutans* cultures exposed to 0.5mM H_2_O_2_. This result is consistent with previous reports (30), and it suggests that PerR is a redox sensor in *S. mutans*. The PerR protein in *Bacillus subtilis* has also been implicated in redox sensing following protein activation (16, 31). Specifically, Fuangthong et al. (2002) describe a role for H_2_O_2_ (or a H_2_O_2_ reaction product) and intracellular Mn^2+^:Fe^2+^ in PerR activation via histidine oxidation (31). While it is possible that a similar mechanism for PerR activation occurs in *S. mutans* given conserved histidine H37 and H101 residues that can undergo metal-catalyzed oxidation, further investigations are warranted to elucidate the details of redox sensing that engage PerR with DNA binding and gene regulation in *S. mutans*.

Unlike *S. mutans* and *S. pyogenes* that harbor a single PerR protein, *B subtilis* harbors two separate Fur family metalloregulators, PerR and Fur (16), the latter sharing 23.1% identity and 42.6% similarity with PerR from *S. mutans* (Fig. S1). An *in silico* analysis of the *S. mutans* genome revealed Fur- and PerR-like binding sequences at numerous putative promoter regions from genes across the UA159 chromosome (Dataset S1), including genes that mediate oxidative stress tolerance via ROS scavenging (eg. *ahpC*, SMU.764) and thiol homeostasis (eg. *gor*, SMU.838). Other binding targets of the PerR homolog were noted in regions upstream of metal ion transport genes (eg. *zupT*, SMU.2069) and *codY*, SMU.1824c), regulators of metal ion homeostasis (*sloR*; SMU.186), and yet other virulence genes that encode sucrose-dependent adherence (*gtfB*; SMU.1004, *ftf*; SMU.2028, *gbpA*; SMU.2112, *gbpC*; SMU.1396, and *gbpD*; SMU.772), acid tolerance (*fabM*; SMU.1746), and two-component signaling systems (*vicR*; SMU.1517). Additional genes preceded by both binding motifs are listed in the supplemental dataset (S1).

The Fur and PerR binding sequences at the *S. mutans sloR* locus share overlap with the −35 and −10 regions of the *sloR* promoter, respectively (Fig. 3), strongly implicating PerR in *S. mutans sloR* gene regulation. In fact, our follow-up investigations of *sloR* transcription in the presence versus absence of PerR support repression of *sloR* by PerR in *S. mutans* cultures challenged with 0.5mM H_2_O_2_, but not in the absence of the peroxide stressor. Moreover, computational analysis did not reveal a SloR recognition element (SRE) proximal to the *perR* promoter (22), consistent with the results of qRT-PCR experiments that indicate no significant impact of the SloR metalloregulator on *perR* gene transcription under conditions of peroxide stress.

The *in silico* prediction of Fur- and PerR-like consensus sequences in the *sloR* promoter, combined with the results of expression profiling experiments that support repression of *sloR* transcription by PerR, led us to hypothesize direct binding of PerR to the *sloR* promoter. To address this hypothesis, we used a heterologous *B. subtilis* Fur protein in gel mobility shift experiments with the *S. mutans sloR* promoter region as the target probe and noted direct Fur binding at protein concentrations as low as 60nM. In addition, Fur binding to a degenerate *S. mutans* probe containing transition mutations in a predicted Fur binding sequence that overlaps the −35 *sloR* promoter region was compromised by 20% when compared to wild type. Taken together, this indicates that the PerR-mediated repression of *sloR* is both direct and sequence-specific. Moreover, Fur appears to bind two sites on the *sloR* promoter probe as indicated by two bands on the gel shift (Fig. 5B). This binding appears to be cooperative since the addition of 15mM EDTA to the reaction mixture abrogated only one of the two band shifts. Taken together, we propose that the PerR protein in *S. mutans* self-associates to form a dimer or higher order oligomer prior to DNA binding at a so-called “Fur-box” in the −35 region of the *sloR* promoter, and at a “PerR box” that is predicted to overlap the-10-promoter region. Such binding would be expected to displace the RNA polymerase holoenzyme, thereby repressing *sloR* gene transcription. A *S. mutans* PerR-DNA interaction may also help maintain low levels of *sloR* transcription in peroxide-challenged *S. mutans* cultures by obstructing positive *sloR* gene control from a promoter-distal SloR binding site described previously (9, 22).

Attempts to shift DNA probes with a recombinant PerR protein that we had prepared from *S. mutans* were unsuccessful, likely due to the absence of cofactors that are needed to facilitate PerR oligomerization and/or -DNA binding *in vitro*. Reports in the literature describe ferrous iron and zinc as necessary cofactors for dimerization and DNA binding of the *B. subtilis* and *S. pyogenes* PerR orthologs (28, 29). The DNA binding activity of PerR is also sensitive to relevant peroxide concentrations which are needed to instigate metal-dependent protein oxidation (32). We reasoned that these and other putative cofactors would be present and available to PerR in protein lysate preparations that derive from cultures exposed to 0.5mM H_2_O_2_. Indeed, the outcome of the gel mobility shift experiments with WCL as the protein source revealed a band shift when the UA159 WCL was allowed to react with a 53-bp *sloR* promoter target sequence (Fig. 7).

Among the antioxidant genes that belong to the *S. mutans* PerR regulon are *tpx* and *dpr* which encode a thiol peroxidase and a ferric iron binding protein, respectively. Although we noted differences in *tpx* and *dpr* transcription that were indirect and relatively modest in the PerR-deficient GMS802 and SloR-deficient GMS584 single knockout mutants compared to wild type (data not shown), their heightened expression was evident in the GMS1386 double mutant. We believe the moderate phenotype of the single mutants speaks to profound redundancies in the PerR and SloR pathways that regulate essential antioxidant gene expression, whereas the more notable phenotype in GMS1386 suggests an interplay between PerR and SloR in *S. mutans* oxidative stress tolerance.

Consistent with resistance of the *perR*-deficient GMS802 strain of *S. mutans* to peroxide stress, and sensitivity of the *sloR*-deficient GMS584 strain to the same stressor (20), we hypothesize a relationship of reciprocity between the PerR and SloR metalloregulators in *S. mutans* cultures challenged with H_2_O_2_. (Fig. 8). Specifically, our experimental findings support a regulatory model that begins with heightened *perR* expression above baseline levels in response to H_2_O_2_ stress, that we believe instigates the metal ion-catalyzed activation of the PerR protein via histidine oxidation (30, 31). Subsequent PerR binding to Fur- and PerR-like consensus sequences is then enabled within the –35 and –10 regions of the *sloR* promoter respectively, which represses *sloR* transcription, causing cytosolic SloR concentrations to fall. Accumulating evidence in our laboratory further indicates that intracellular SloR concentrations are subject to post-translational degradation by the *S. mutans* Clp proteolytic system (unpublished observations) under conditions of peroxide stress. Hence, a decline in cytosolic SloR levels in the face of peroxide challenge compromises the positive influence that SloR would otherwise have on the *S. mutans* oxidative stress response. The concomitant rise in PerR activity further exacerbates downregulation of the oxidative stress response, in part by repressing transcription of the antioxidant genes *dpr* and *tpx*, albeit indirectly. Consistent with these findings are reports implicating the *S. mutans* PerR regulator as a repressor of *dpr* and *sod* transcription (27). In addition, a recent report describes co-repression of the *S. mutans spxA1* gene by the PerR and SloR metalloregulators (33). The *spxA1* gene product has been previously described as a positive regulator of oxidative stress tolerance in *S. mutans* (33–35). Clearly, further investigation is warranted to decipher the complexities of antioxidant gene regulation involving SpxA1.

**Fig. 8.**
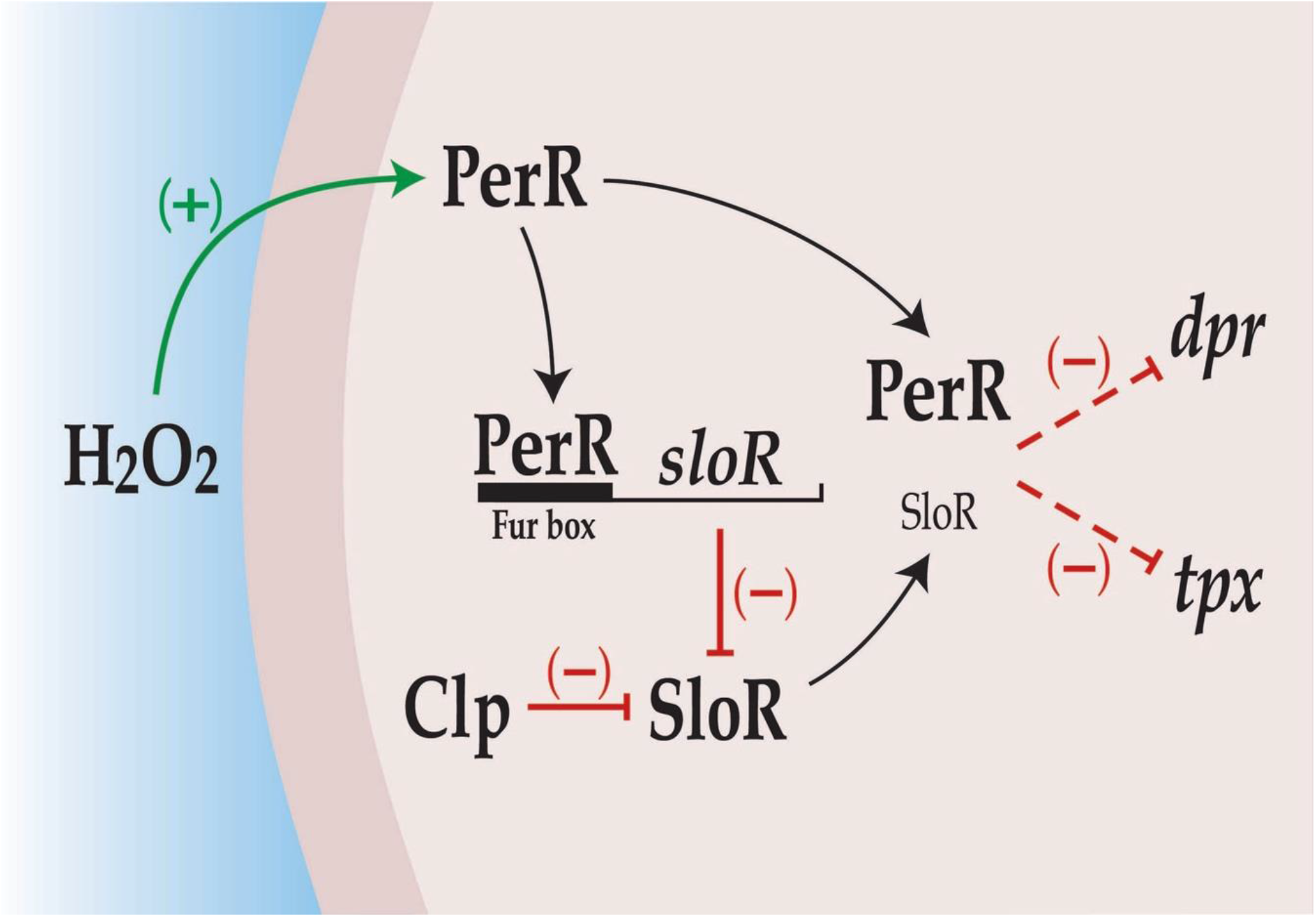
Model of PerR and SloR involvement in the *S. mutans* oxidative stress response. H_2_O_2_ stress and redox sensing by baseline levels of cytosolic PerR upregulate *S. mutans perR* gene transcription. As intracellular concentrations of the *perR* gene product accumulate, PerR binds directly to the *sloR* promoter at conserved Fur- and PerR-like consensus sequences (GATAATAAAACTTTTATC and ATATAATGATTTTGA, respectively) to repress *sloR* gene transcription. SloR is also subject to posttranslational degradation by the *S. mutans* Clp proteolytic system under conditions of peroxide stress. The accumulation of PerR in the cell combined with decreasing intracellular SloR concentrations repress transcription of the *S. mutans* antioxidant genes, *dpr* and *tpx* consistent with the proposed involvement of both SloR and PerR in *S. mutans* stress adaptation.

In summary, we propose that PerR is a prevalent redox sensing metalloregulator in *S. mutans* and that H_2_O_2_ is the possible signaling molecule that activates PerR and promotes *S. mutans* stress adaptation. In the present study, we describe stress adaptation that involves PerR-mediated repression of the *S. mutans dpr* and *tpx* antioxidant genes under conditions of oxidative stress, as well as repression of *sloR* gene transcription. Why the *S. mutans* PerR metalloregulator evolved to repress expression of its antioxidant arsenal in the face of peroxide stress is an interesting question. We propose the answer resides in the two faces of reactive oxygen species (ROS) – redox biology which activates signaling pathways to initiate normal biological processes (36, 37), and oxidative stress when elevated levels of ROS can incur damage to cellular DNA, protein, and/or lipids (13, 15). Taken together with our experimental findings, we hypothesize that PerR repression and the regulatory mechanisms that maintain low levels of SloR in the cell cytosol likely evolved to maintain beneficial redox biology in the face of H_2_O_2_ challenge. In contrast, mobilizers of *S. mutans dpr* and *tpx* transcription, which may involve SpxA1 as a positive regulator, likely evolved as part of an oxidative stress response to prevent ROS-induced cell death. Taken collectively, continuing investigations of the mechanisms that repress the oxidative stress response in *S. mutans* combined with the regulatory events that facilitate oxidative stress tolerance can elucidate *S. mutans* fitness and survival strategies in healthy versus dysbiotic plaque, and so inform an anti-caries therapeutic.

## MATERIALS AND METHODS

### Bacterial strains, plasmids, and primers

The bacterial strains and plasmids that were used in this study are presented in Table S1. Primers and oligonucleotides are presented in Table S2. Primers were designed using MacVector 7.0 software and purchased from Eurofins Genomics (Louisville, KY).

### Bacteriological media and reagents

*S. mutans* strains UA159, GMS584, GMS802, and GMS1386 were grown as standing cultures at 37°C and 5% CO_2_ in 15-mL Falcon tubes (Fisher Scientific, Pittsburgh, PA) in Todd-Hewitt broth supplemented with 0.3% yeast extract (THYE). Erythromycin (10 μg/ml) was added to THYE broth when growing *S. mutans* GMS584 and GMS802 cultures. Erythromycin (10 μg/ml) and kanamycin (250μg/mL) were added to THYE broth when growing *S. mutans* GMS1386.

*S. gordonii* Challis CH1 cells were propagated in THYE at 37°C with gentle aeration to achieve “aerobic” growth. The ‘conditioned medium’ used in growth rate determination assays was formulated as previously described (12) with the following modifications: first, *S. gordonii* grown overnight in THYE broth was used to inoculate 40 mL of prewarmed THYE to an optical density (OD_600nm_) of 0.025. The subculture was then grown with aeration on a shaking platform (150 rpm) to mid-logarithmic phase (OD_600nm_ ~ 0.6) after which the cells were harvested by centrifugation at 1,700 x g for 10 minutes in a Sorvall SS34 rotor. 20mL of the resulting culture supernatant was then supplemented with 0.25% glucose and filter sterilized. The culture supernatant is referred to as “conditioned medium” in subsequent growth determination assays.

### Construction of *S. mutans perR* insertion-deletion mutants

A 980-bp amplicon harboring the 477-bp *perR* coding sequence (SMU.593) along with its upstream promoter region, was amplified with primers 593.smaI.F and 593.smaI.R (Table S2), digested with *Sma*I (Promega), and ligated into a unique *Sma*I restriction site on the pBGK2 integration vector (38). The resulting ligation mixture was introduced into chemically competent *E. coli* GC5 by natural transformation in accordance with the instructions of the manufacturer (Gene Choice), and transformants were selected on L-agar supplemented with kanamycin. Plasmid DNA was isolated from select transformants using a miniprep kit according to the instructions of the manufacturer (QIAGEN), and the recombinant construct, called pJS2, was confirmed via restriction mapping.

PCR ligation mutagenesis (39) was then performed to disrupt the *perR* gene on the wild type *S. mutans* UA159 chromosome. Specifically, primers 593.P1/ 593.P2, and 593.P3/593.P4 (Table S2) were used to amplify the 5’ and 3’ flanking regions of the SMU.593 gene from the *S. mutans* genome, thereby generating a 103-bp deletion in the *perR* coding sequence. The resulting P1/P2 and P3/P4 amplicons were digested with *Asc*I and *FseI* (New England Biolabs) and ligated in separate bimolecular reactions to an *ermAM* erythromycin resistance cassette with compatible 5’ *Asc*I and 3’ *Fse*I overhangs. The ligation mixtures were then used as a template for a second round of PCR (as described above) with primers 593.P1 and 593.P4 (Table S2), to generate a P1/P2-ermAM-P3/P4 tripartite construct. The final amplification mixture was used to transform *S. mutans* UA159 in the presence of competence-stimulating peptide (CSP) as described previously (40). Candidate transformants were screened for erythromycin resistance on THYE-erm agar, and disruption of the SMU.593 coding sequence was confirmed by PCR amplification and sequencing of the *perR* locus using primers 593.P1 and 593.P4. The resulting insertion-deletion mutant was named GMS802.

The pJS2 integration construct was used to complement the SMU.593 mutation in *S. mutans* GMS802 *in cis*. The recombinant plasmid was introduced into the *S. mutans* UA159 genome in single copy by CSP-induced transformation as previously described (40). The pBGK2 parent vector was used to transform GMS802 in parallel as a negative control. Successful complementation of the SMU.593-specific mutation in GMS802 was confirmed phenotypically in *S. gordonii* antagonism assays, and the complemented *S. mutans* strain was named GMS803.

To generate a double *perR* and *sloR* insertion-deletion mutant, an allelic exchange strategy similar to that described for GMS802 was employed to disrupt the *perR* gene in the *S. mutans* SloR-deficient GMS584 strain generated previously in our laboratory (21). A 1500-bp *aphA3* cassette was inserted into the *perR* coding sequence, and the resulting insertion-deletion mutant, called GMS1386, was confirmed by PCR and nucleotide sequencing across the *perR-aphA3* and *sloR-ermAM* junctions.

### Growth determination assays

*S. mutans* UA159 and GMS802 were monitored for growth in a Bioscreen C microbiology plate reader (Growth Curves, U.S.A.) as previously described (21). To assess peroxide tolerance, mid-exponential phase cells (OD_600nm_ 0.4-0.6) were diluted 1:50 in THYE containing either 0% H_2_O_2_ or 0.0015% H_2_O_2_, or else in a ‘conditioned medium’ prepared as described above (12). The Bioscreen reader was programmed with Biolink software (Labsystems) to monitor growth at 37°C for 24 h with low amplitude shaking every 15 min. To assess the growth of UA159 and GMS802 under aerobic conditions, the Bioscreen plate reader was programmed for continuous low amplitude shaking and monitored over a 24 h period. Generation times were calculated using the equation dt = ln(2)/(15/(OD_600_B-OD_600_A)). The resulting data were interpreted by applying Univariate ANOVA and post-hoc Bonferrioni T-tests with SPSS statistical analysis software (SPSS for Windows).

### Hydrogen peroxide challenge in broth culture

*S. mutans* cultures were grown as described above in THYE broth to an OD_600_ of 0.5 (±0.05) as determined by spectrophotometry (GENESYS 20, Thermo Scientific). 0.5mM H_2_O_2_ was added to select broth cultures for a cell exposure time of fifteen minutes, after which the cells were immediately harvested by centrifugation and lysed for nucleic acid isolation.

### Antagonism assays

The sensitivity of *S. mutans* UA159 and GMS802 to H_2_O_2_ antagonism by *S. gordonii* was assessed in plate assays as described previously (12) with the following modifications. 8 μl of an overnight culture of *S. gordonii* was spot inoculated onto a THYE agar plate and incubated at 37°C and 5% CO_2_ for 16 hours to allow for production of hydrogen peroxide, after which 8 μl of a *S. mutans* UA159, GMS802, or GMS803 culture was spot inoculated proximal to *S. gordonii*. Plates were incubated as described above for an additional 14 hours and the resulting zones of growth and growth inhibition were qualitatively assessed.

### *In silico* prediction of *S. mutans* PerR and Fur binding sites

The CollecTF database of bacterial transcription factor binding sites (41) was used to identify consensus sequences for Fur homologs (including PerR) in *Bacillus sp*. TRANSFAC position weight matrices for each consensus sequence were converted to MEME format and used to search extracted intergenic regions with a minimum length of 50-bp from the *S. mutans* UA159 genome using FIMO version 5.3.0 (42) at the MEME-suite webserver (43) using a p-value threshold of 10^-3^ and a bootstrap implementation (44) to estimate false discovery rates as q-values (45). Genes bordering intergenic regions were identified and distances of motif matches from start codons were calculated. Matches with the closest boundary within 50-bp of the 5’ end of a neighboring gene were retained and then filtered with the lowest p-value motif match reserved for each matrix search.

### RNA isolation and cDNA synthesis

Total RNA was isolated from mid-logarithmic *S. mutans* cultures grown in THYE as described previously (46). RNA was assessed for integrity on a 0.8% agarose gel, adjusted to a concentration of 100 ng/μl using a NanoDrop Lite spectrophotometer (Thermo Fisher Scientific, Waltham, MA), and immediately reverse transcribed into cDNA using a Revert-Aid First-Strand cDNA Synthesis Kit (Thermo-Scientific) according to the manufacturer’s instructions. Reaction mixtures with nuclease-free water instead of RNA served as no-template controls, and reactions without reverse transcriptase were prepared as RT^-^ controls. cDNA integrity was examined on a 0.8% agarose gel and compared to the RT^-^ controls.

### Semi-quantitative real-time PCR (qRT-PCR)

qRT-PCR was performed with cDNA generated as described above. Amplification was performed in a CFX96 real-time system (Bio-Rad) with SYBR Green as the intercalating dye (Bio-Rad) and 500nM *perR, sloR, dpr, tpx*, and *hk11*-specific primers (**Table S2**). RT^-^ controls were run in parallel with the same primers for each cDNA template to confirm the absence of contaminating genomic DNA. The thermal cycling conditions were 95°C for 3 min followed by 40 cycles of 95°C for 10 s, 60°C for 10 s, and 72°C for 15 s. Primer efficiency was calculated using a standard curve using dilutions of UA159 cDNA. Gene expression was normalized against the expression of the endogenous control gene *hk11*, the expression of which did not change under the experimental test conditions.

### *S. mutans* whole cell lysate preparation

*S. mutans* UA159 were grown to mid-exponential phase (OD_600nm_ 0.5) in 30 mL of prewarmed THYE broth and with selective pressure when appropriate. The cells were harvested by centrifugation at 8,000 × *g* for 10 min in a FiberLite F13S-14x50cy rotor and then washed with 30 ml of 10 mM Tris-Cl, pH 7.8. Cells were resuspended in 750 μl of 10 mM Tris-Cl, pH 7.8, transferred to an ice-cold 2-ml screw-cap tube containing 300 μl of 0.1-mm zirconium beads, and mechanically disrupted at 4°C for 30 s at a speed of 5.0 in a BIO 101 FastPrep machine. The tubes were then centrifuged at 18,000 × *g* for 10 min at 4°C, and wholecell lysates were collected as supernatants and stored at −20°C.

### Preparation of DNA target probes for gel mobility shift experiments

Q5 High-Fidelity DNA Polymerase (New England BioLabs) was used to amplify target probes from the *S. mutans* UA159 chromosome with *sloR*- and *tpx*-specific primers (Table S2) according to the recommendations of the supplier. The NEBtools Tm calculator was used to facilitate primer design and define appropriate annealing temperatures. PCR products were confirmed on a 0.8% agarose gel and visualized after staining with SYBR Safe DNA Gel Stain (Invitrogen, Carlsbad, CA). The amplicons were purified with an Invitrogen PCR Purification Kit (ThermoFisher, Waltham, MA) according to the manufacturer’s instructions prior to end-labeling.

Wild type and degenerate versions of the predicted PerR binding region in the *S. mutans sloR* promoter region were prepared as oligonucleotides (Table S2) and purchased from Eurofins Genomics (Louisville, KY). The forward (5’ to 3’) oligo was annealed to its reverse complement in 10X annealing buffer containing 100mM potassium acetate and 30μM HEPES at pH 7.5. The annealing conditions were set in a thermal cycler at 95°C for 2 minutes, followed by -0.1°C every cycle for 450 cycles, and then held at 4°C.

### Gel mobility shift assay

Gel mobility shift assays were performed as previously described (20, 23). Purified PCR products or annealed oligonucleotides were prepared as described above and end-labeled with 10 μCi [γ-^32^P]ATP (Perkin-Elmer) and 10 U T4 polynucleotide kinase according to established protocols (New England BioLabs). Binding reactions included 1 μl (~30 ng) of end-labeled DNA and up to 400nM *B. subtilis* Fur or 6.5ug of UA159 or GMS802 whole cell lysate in a total reaction volume of 16uL Reaction mixtures without a source of protein were included as negative controls. All samples were resolved on 12% nondenaturing polyacrylamide gels for up to 70 min at 300 V. Gels were exposed to Carestream BIOMAX XAR Film and stored at −80°C for up to 48 hours in the presence of an intensifying screen. Autoradiography followed in the dark room according to established protocols. ImageJ software, version 2.0.0 (National Institute of Health) was used to quantify the band shift intensities on autoradiograms. The band shift, expressed as a percentage, was calculated by comparing the mean pixel area of the 60nM Fur band shift with that of the unshifted target sequence in the same lane, and then expressed as a percentage of the 100% band shift that we observed when 400nM Fur was used to shift the target probe.

## SUPPLEMENTAL MATERIAL

**Table S1.**
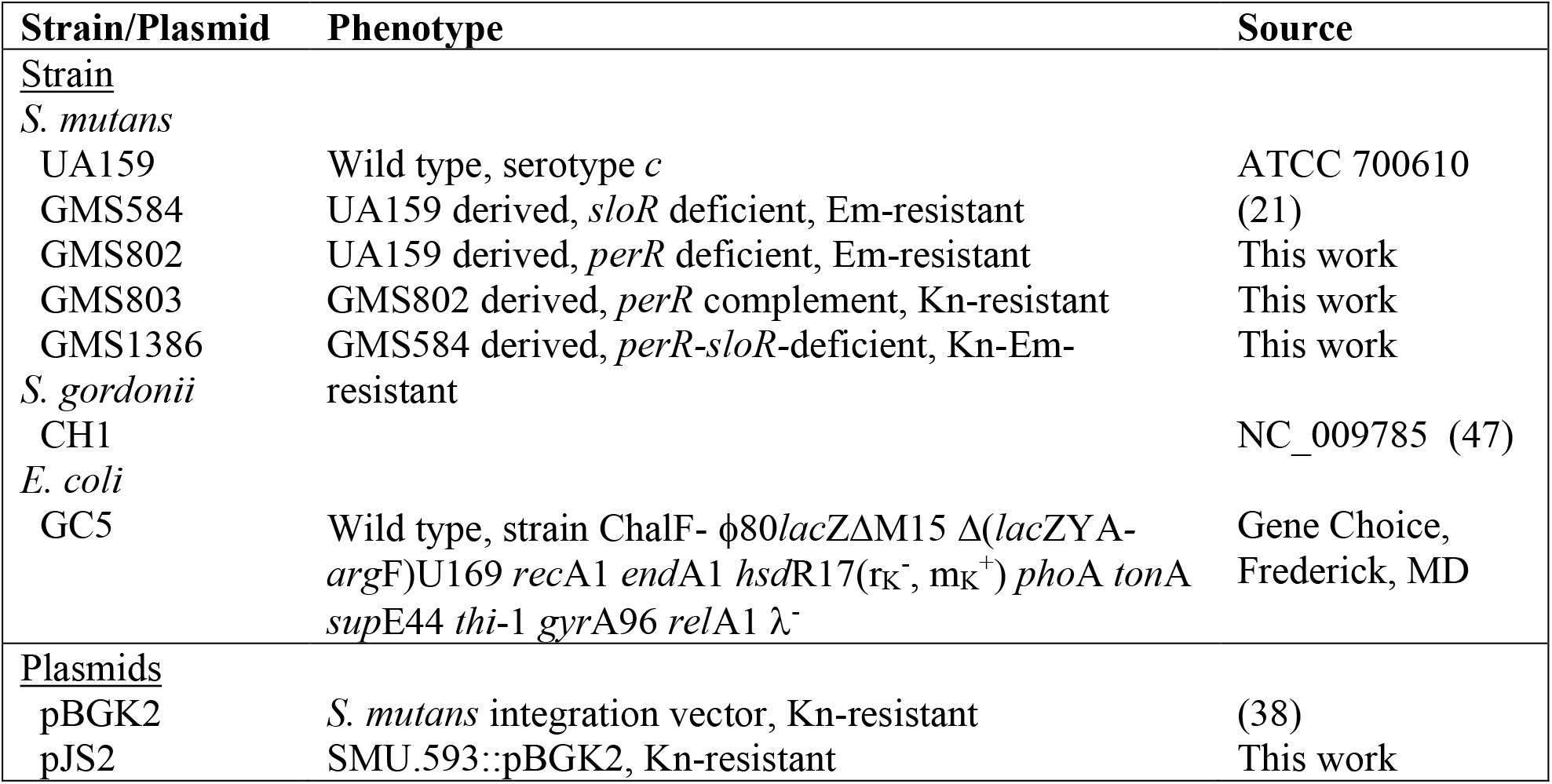
Bacterial strains and plasmids used in this study.

| Strain/Plasmid | Phenotype | Source |
| --- | --- | --- |
| <u>Strain</u> |  |  |
| *S. mutans* |  |  |
| UA159 | Wild type, serotype c | ATCC 700610 |
| GMS584 | UA159 derived, *sloR* deficient, Em-resistant | (21) |
| GMS802 | UA159 derived, *perR* deficient, Em-resistant | This work |
| GMS803 | GMS802 derived, *perR* complement, Kn-resistant | This work |
| GMS1386 | GMS584 derived, *perR-sloR*-deficient, Kn-Em-resistant | This work |
| *S. gordonii* |  |  |
| CH1 |  | NC_009785 (47) |
| *E. coli* |  |  |
| GC5 | Wild type, strain ChalF- $\phi 80lacZ\Delta M15 \Delta(lacZYA-argF)U169 recA1 endA1 hsdR17(r_K^-, m_K^+) phoA tonA supE44 thi-1 gyrA96 relA1 \lambda^-$ | Gene Choice, Frederick, MD |
| <u>Plasmids</u> |  |  |
| pBGK2 | *S. mutans* integration vector, Kn-resistant | (38) |
| pJS2 | SMU.593::pBGK2, Kn-resistant | This work |

**Table S2.**
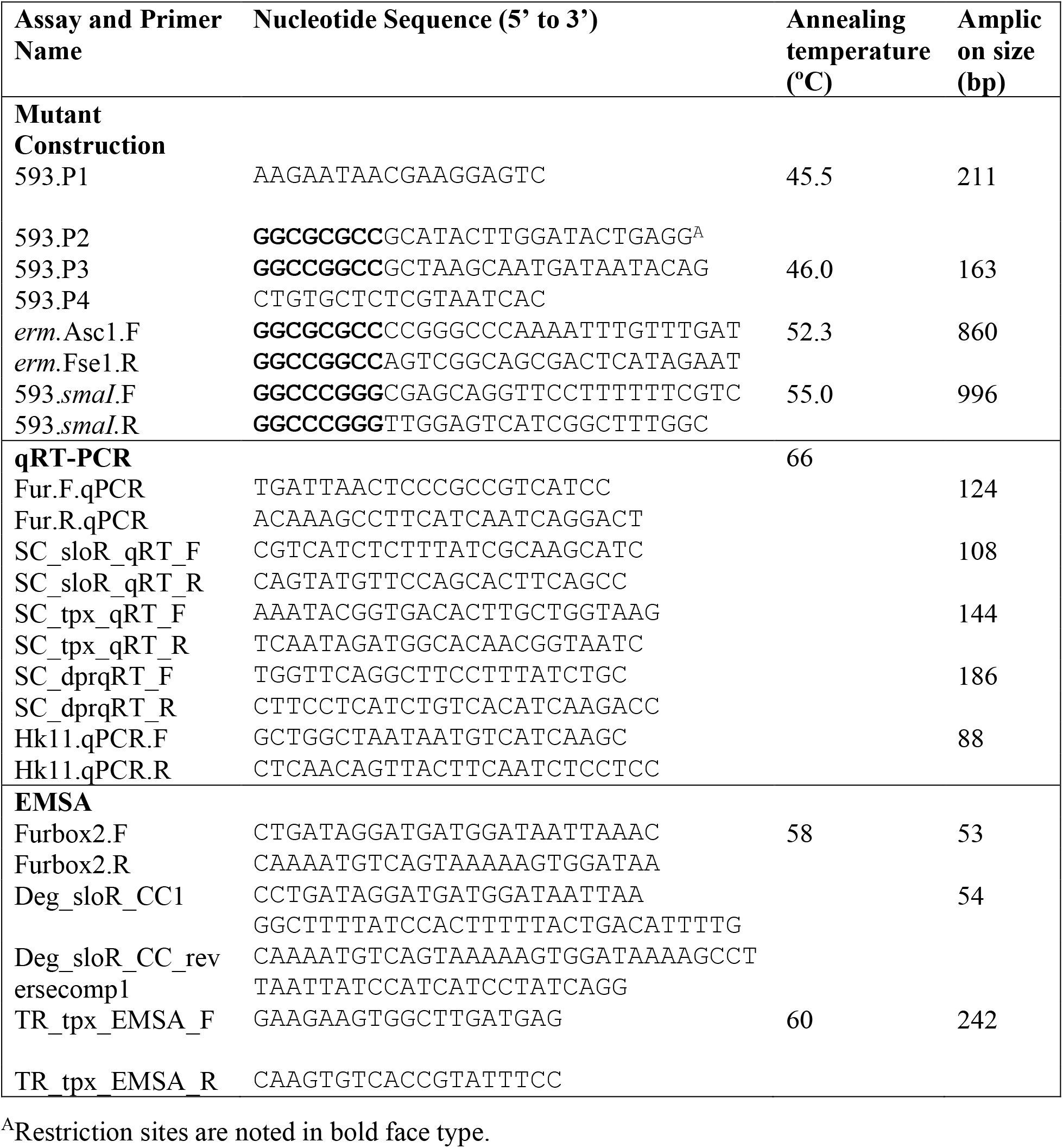
Primers and oligonucleotides used in this study.

**S1.**
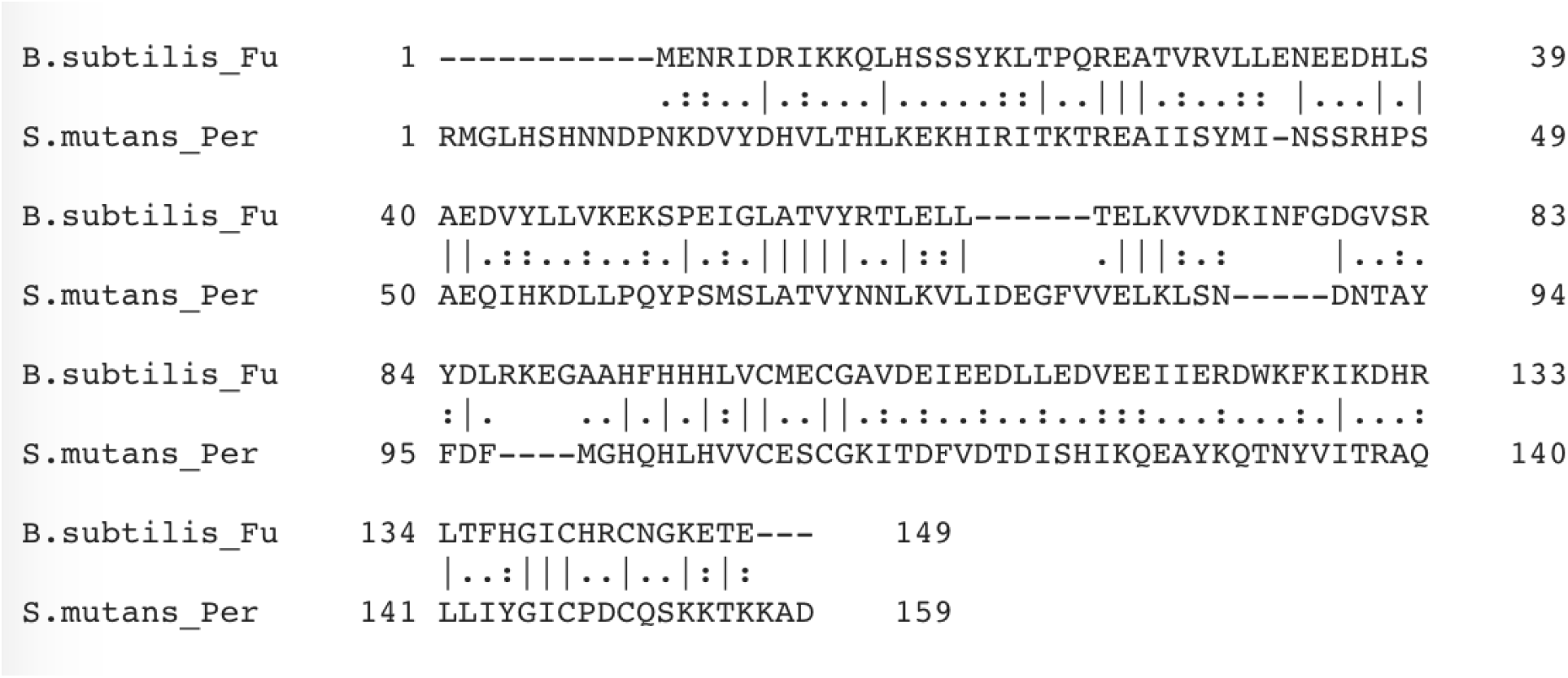
Alignment of the *B. subtilis* Fur and *S. mutans* PerR primary amino acid sequences. Shown is a local sequence alignment of the 17.4kDa Fur and 18kDa PerR protein sequences from *B. subtilis* and *S. mutans*, respectively, using EMBOSS Matcher (EMBL-EBI). Amino acid sequence identity and similarity are indicated by vertical dashes (|). Conserved amino acid substitutions are marked with a colon (:) if they are strongly similar and have a score > 0.5 in a PAM250 matrix, or with a period (.) if they are only weakly similar, scoring ≤ 0.5 in a PAM250 matrix. The *S. mutans* PerR protein shares 23.1% amino acid sequence identity and 42.6% similarity with the Fur protein from *B. subtilis*.

## ACKNOWLEDGEMENTS

This research was supported by NIH grant DE014711 to G.S., Undergraduate Research Fellowships from the American Society for Microbiology and the Vermont Genetics Network to J.S., and by the Middlebury College Department of Biology. We acknowledge Gary Nelson for figure preparation and Dr. John Helmann for providing us with Fur protein from *Bacillus subtilis*. We declare that we have no conflicts of interest to report.

